# Synergistic stabilization of a double mutant in CI2 from an *in-cell* library screen

**DOI:** 10.1101/2020.12.01.406082

**Authors:** Louise Hamborg, Daniele Granata, Johan G. Olsen, Jennifer Virginia Roche, Lasse Ebdrup Pedersen, Alex Toftgaard Nielsen, Kresten Lindorff-Larsen, Kaare Teilum

## Abstract

Most single point mutations destabilize folded proteins. Mutations that stabilize a protein typically only have a small effect and multiple mutations are often needed to substantially increase the stability. Multiple point mutations may act synergistically on the stability, and it is not straightforward to predict their combined effect from the individual contributions. Here, we have applied an efficient *in-cell* assay to select variants of the barley chymotrypsin inhibitor 2 with increased stability. We find two variants that are more than 3.8 kJ/mol more stable than the wild-type. In one case the increased stability is the effect of the single substitution D55G. The other case is a double mutant, L49I/I57V, which is 5.1 kJ/mol more stable than the sum of the effects of the individual mutations. In addition to demonstrating the strength of our selection system for finding stabilizing mutations, our work also demonstrate how subtle conformational effects may modulate stability.

## Introduction

Understanding how the stability of a protein changes when an amino acid residue is changed is fundamental for several biological processes and the aetiology of many diseases (Stein et al., 2019). For using proteins in biotechnological and biopharmaceutical applications it is often an advantage that the proteins have long shelf-lives and are not degraded too rapidly during the applications (Modarres et al., 2016). Our ability to engineer proteins with increased stability or to understand how amino acid changes cause decreased stability has thus been the subject of large number of studies.

We recently described a system based on recombinant expression in *E. coli*, that can be used to measure both protein translation and folding stability *in vivo* (Figure 1a) (Zutz et al., 2020). The translation sensor is based on an RNA hairpin structure inserted into a polycistronic mRNA coding for the protein of interest and for the fluorescent protein mCherry, thus making it possible to read out efficient translation through a red fluorescent signal. The protein folding and stability sensor is based on GFP-ASV, an unstable GFP variant, through a system engineered to be expressed as a response to protein misfolding, and the green fluorescence is used as a proxy for *in vivo* protein stability. This misfolding response relies on a heat shock promoter, lbpAp, and the *E. coli* heat shock system. With increasing levels of protein misfolding, more of the chaperone DnaK will bind the misfolded protein instead of the *E. coli* heat shock sigma factor, RpoH. In the absence of DnaK, RpoH can participate in assembly of the RNA polymerase sigma 32 complex, which can drive transcription from the lbpAp promoter. The presence of misfolded protein that can bind DnaK thus results in the expression of GFP and a green fluorescence signal. By expressing libraries of random mutations in a given protein in this bacterial sensor system and analysing the cells by fluorescence-activated cell sorting (FACS), it is possible to select large sets of protein variants that retain a folded structure, thus avoiding complications from using a functional assay as a proxy for folding stability.

**Figure 1.**
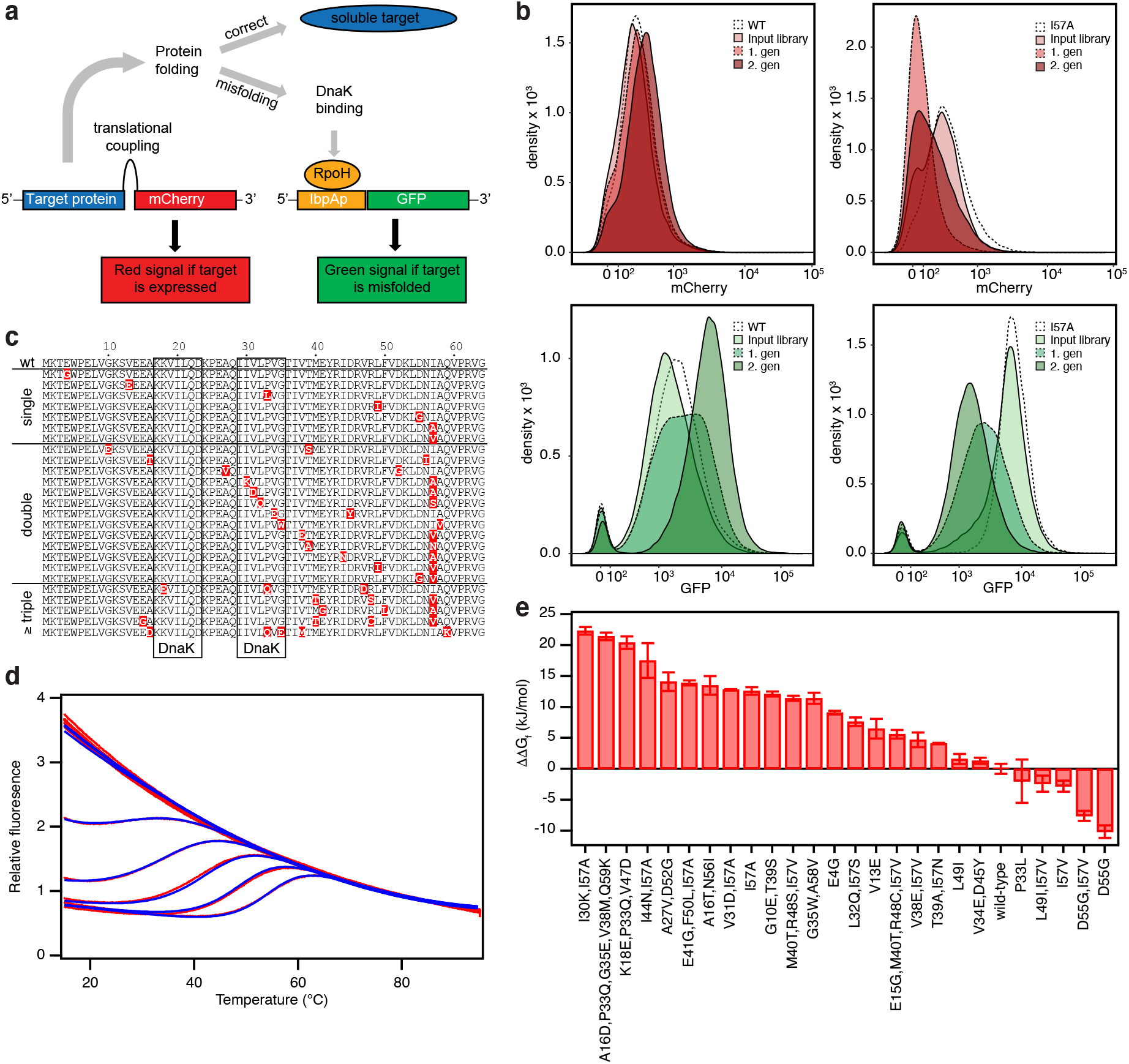
Selection and stability of CI2 variants. (**a**) Overview of the bacterial folding sensor. The expression of mCherry is linked to the expression of the target protein. Cells expressing a target protein will thus also have red fluorescence. The expression of GFP is linked to the presence of misfolded protein. Cells expression a target protein with high tendency of misfolding will thus have a high level of green fluorescence. (**b**) FACS profiles of *E. coli* cultures expressing libraries of random mutations in wildtype CI2 (left column) and in CI2,I57A (right column). mCherry fluorescence and GPF fluorescence are shown in the upper and lower rows, respectively. Profiles for the background variants of CI2, the input mutant library and the two rounds of sorting are shown. (**c**) Sequence alignment of 26 CI2 variants purified and showing two-state behaviour in equilibrium stability measurements. (**d**) Thermal unfolding curves of CI2,I57A measured by fluorescence at 350 nm at 13 concentrations of GuHCl ranging from 0 to 5 M. The experimental data are shown in red and the fits to a model for two-state folding as blue lines. (**e**) Difference in conformational stability, ΔΔ*G*_f_, relative to wild-type CI2 for 26 variants. The error bars are the standard deviation on Δ*G*_f_ listed in Table 1 that was propagated from the standard deviations on *T*_m_, Δ*H*_m_ and Δ*C*_p_ obtained from the non-linear global fits of the fluorescence unfolding data.

An alternative to screening mutant libraries for proteins with altered stability is to calculate the effect of substituting amino acids and find variants with the desired properties. Several computational tools have been developed that predict the change in free energy for folding (ΔΔ*G*_f_) between a wild-type protein and a mutant (Capriotti et al., 2005; Dehouck et al., 2009; Goldenzweig et al., 2018; Guerois et al., 2002; Jochens et al., 2010; Kaufmann et al., 2010; Kellogg et al., 2010; Pandurangan et al., 2017; Parthiban et al., 2006; Reetz et al., 2006; Rohl et al., 2004; Schymkowitz et al., 2005; Steipe et al., 1994; Sullivan et al., 2012; Trudeau et al., 2014; Wijma et al., 2014; Yamashiro et al., 2010). In general, the methods perform rather well when predicting the effects of destabilizing mutations but often fail in predicting stabilizing mutations (Foit et al., 2009). A comparison of several stability predictors showed an average correlation of around 0.6 between experimentally determined and computed changes in stability for all types of mutations (Khan and Vihinen, 2010; Potapov et al., 2009). The algorithms are better at predicting deletion mutations in the hydrophobic core, than mutations that increase the size of the side chain, mutations on the protein surface and mutations where electrostatic interactions contribute to the stabilization. This is partly a result of the data available for training the algorithms that mainly consist of deletion mutations in the hydrophobic core (Gromiha et al., 2016). A particular challenge in predicting stabilizing protein variants is that among the few single substitutions that are actually stabilizing the effects are often small, so that multiple substitutions may be needed to create a substantial stabilizing effect (Goldenzweig et al., 2018). As the effects of the mutations are not always independent, and non-additivity may result in both positive or negative epistasis (Bershtein et al., 2006; Sarkisyan et al., 2016), it can be difficult to predict the stability of proteins with multiple substitutions. One way to improve the computational methods is to generate stability data on a larger set of protein variants generated to scan sequence space better than the current available datasets and including also stabilizing variants.

Here, we have applied the bacterial sensor to select variants from a library of random mutations of barley chymotrypsin inhibitor 2 (CI2) to broadly cover sequence and stability space. CI2 is a small single domain protein of 64 residues, which has been extensively used as a model to understand key concepts of protein folding and stability (Itzhaki et al., 1995; Jackson et al., 1993; Jackson and Fersht, 1991a, 1991b; Neira et al., 1997). Our aim in the current work was two-fold. First, we wanted to demonstrate how the sensor system can be used to select for proteins with increased stability. Second, we aimed to generate a set of data for benchmarking and later optimization of stability predictors. While many of the 25 variants for which we measured the stability, destabilize CI2, we found two variants, L49I/I57V and D55G, that are significantly stabilized relative to wild-type CI2. For L49I/I57V there is a strong positive synergistic effect between the two substitutions and this variant is stabilized by 5.1 kJ/mol more than the sum of the individual effects of the two single variants. A detailed analysis of the structural changes in L49I/I57V suggests that several subtle long-range effects underlie the high stability gain.

## Results

### FACS sorting libraries of random CI2 variants

CI2 is a highly stable protein with free energy for folding, Δ*G*_f_ = 31 kJ/mol (Hamborg et al., 2020; Itzhaki et al., 1995) and, as previously shown (Zutz et al., 2020), when wild-type CI2 is expressed in the bacterial sensor system (Figure 1a) only little GFP is produced (Figure 1b). The dynamic range of the GFP signal towards discovering stabilized variants (i.e. less GFP) is thus very small. A library of random mutations in wild-type CI2 will thus be most suited for selecting variants with stabilities that are lower than the wild-type protein but that are still able to fold. To select variants of CI2 that are more stable than the starting point we therefore opted to use a destabilized background as starting point. The I57A variant of CI2 is significantly destabilized (Δ*G*_f_ = 14 kJ/mol) and results in a high GFP signal in the sensor system (Figure 1b). With a library of random mutations in I57A there will be a large dynamic range in the GFP signal towards more stable variants with less GFP. This library will thus be suited for selecting variants of CI2 with stabilities higher than I57A. Consequently, we prepared two libraries of random mutations with expected mutation frequencies of 0-4 amino acid residues per gene in the background of the wild-type sequence and of the I57A sequence, respectively.

The mutant libraries that had sizes of 16000 – 80000 were expressed in the dual-sensor system in *E. coli* and analysed by FACS after one hour of induced protein expression. As expected, the GFP fluorescence observed in the initial FACS run of the library made from wild-type CI2 is low and similar to that seen for an *E. coli* culture expressing non-mutagenized wild-type CI2 in the sensor system (Figure 1b). To screen for destabilized protein variants in this library, cells were sorted for high GFP fluorescence, defined as the upper 1-10 % of the GFP signal. The selected cells were grown and sorted twice more using the same criteria (Figure 1b). After each round of sorting, a clear shift in the GFP fluorescence is observed corresponding to an enrichment of clones expressing CI2 variants with decreased stability compared to the wild-type protein. In the same way, the library made in the I57A background was repeatedly sorted for the lower 1-10 % of the GFP signal, resulting in a clear shift from high to low GFP fluorescence (Figure 1b) and enrichment of the library with clones expressing CI2 variants that activate the misfolding sensor less compared to I57A.

### Selecting variants for further analysis

To identify the protein variants in the two final libraries we randomly selected 118 clones from the library starting from the wild-type sequence and 289 clones from the library starting from the I57A sequence by collection of single cells from the last rounds of FACS screening. These clones were characterised by Sanger sequencing. In addition, we also sequenced the final I57A library by next-generation sequencing (NGS). In total we found 71 unique sequences without stop codons, deletions or insertions in the region encoding the 64 amino acid residues of CI2. 41 of the sequences were from the wild-type library and 30 sequences were from the I57A library (Figure S1).

To express and purify the 71 CI2 variants, they were subcloned into pET11a without the hexa-His tag, which is part of the folding sensor system. The presence of this C-terminal His-tag interferes with key interactions of the CI2 C-terminal carboxylate and compromises the stability of the protein during purification. However, the destabilization gives just the right stability window for finding variants with altered stability in the sensor system. Thus, the FACS selection was done on CI2 libraries with the His-tag, whereas the *in vitro* stability measurements were performed on CI2 variants without the His-tag.

Although we selected for protein variants that do not activate the misfolding sensor, some of the variants still did not behave well in the expression system and did not result in pure protein. Of the remaining variants, some did not give useful data in the equilibrium unfolding experiment due to aggregation or multi-state behaviour. We thus ended with 13 unique variants in the wild-type background and 12 unique variants in the I57A background that could be used for stability measurements (Figure 1c). In the set of variants that we have analysed we also included L49I and D55G (*vide infra*).

We note that a large proportion of the variants that were not included in the final set included substitutions in the DnaK binding sites predicted at positions 17-23 and 29-35 (Durme et al., 2009), with the second predicted to be the strongest (Figure S2a). As these variants are expected to interact less well with DnaK, less GFP will also be expressed and they will have the same signature of GFP fluorescence as variants with increased stability even if they are severely destabilized. Examples include substitutions at positions 29 (I29N, I29T), 30 (I30K), 31 (V31D, V31G) and 34 (V34E) that are all predicted to decrease DnaK binding (Figure S2b), and several of these were found in multiple of the variants that could not be purified.

To measure the *in vitro* stability for folding we used our recently described combined two-dimensional thermal and chemical protein unfolding assay, where the unfolding of the protein is followed by the change in intrinsic Trp fluorescence as the temperature is increased at multiple concentrations of denaturant (Figure 1d) (Hamborg et al., 2020). The stabilities of the 27 CI2 variants cover a broad range from −7.4 kJ/mol to −38.5 kJ/mol including five variants that are more stable than the wild-type protein (Table 1 and Figure 1e). We find a strong correlation between Δ*G*_f_ at 25°C and the melting temperature, *T*_m_, which may thus be used as an additional parameter for comparing the stability. Except P33L, all variants selected in the wild-type background are destabilized and scattered throughout the sequence. P33L is stabilized by 1.5 kJ/mol but aggregates when the temperature is above 50 °C unless [GuHCl] > 2 M. All variants selected in the I57A background that are more stable than this background have a valine at position 57; we note that our starting point was designed to avoid random reversion to isoleucine by single nucleotide mutations (see Discussion). From previous work it is known that I57V is slightly more stable than the wild-type protein (Itzhaki et al., 1995), and valine at position 57 is also preferred in CI2 from many other plant species (Lawrence et al., 2010). Most other mutations that occur together with I57V are less stable than I57V alone, but still more stable than the I57A background. There are however two exceptions. Both the L49I/I57V and D55G/I57V are even further stabilized than I57V. Both I at position 49 and G at position 55 are often seen in CI2 from other species (Lawrence et al., 2010). The L49I and D55G variants have not previously been characterized so we also included these single variants in our analysis.

**Table 1.** Thermodynamic data for purified variants of CI2 identified with the folding sensor.

| Variant | $T_m$ (K) | $\Delta H_m$ (kJ/mol) | $\Delta C_p$ (kJ/mol/K) | $m$ (kJ/mol/M) | $\Delta G_f$ (kJ/mol) | $[D]_{50\%}$ (M) |
| --- | --- | --- | --- | --- | --- | --- |
| #I30K, I57A | 327.4 ± 3.1 | 99 ± 11 | 0.4 ± 0.8 | 8.8 ± 1.8 | -8.2 ± 0.6 | 0.9 ± 0.2 |
| A16D, P33Q, G35E, V38M, Q59K | 333.3 ± 1.9 | 134 ± 9 | 2.6 ± 0.3 | 5.4 ± 0.5 | -9.1 ± 0.6 | 1.7 ± 0.2 |
| K18E, P33Q, V47D | 336.0 ± 1.7 | 147 ± 10 | 2.9 ± 0.1 | 4.3 ± 0.2 | -10.1 ± 1.0 | 2.3 ± 0.2 |
| #I44N, I57A | 339.0 ± 2.4 | 163 ± 28 | 2.6 ± 0.4 | 7.4 ± 1.5 | -13.0 ± 2.8 | 1.8 ± 0.5 |
| A27V, D52G | 341.7 ± 1.0 | 211 ± 19 | 3.6 ± 0.3 | 6.0 ± 0.8 | -16.4 ± 1.5 | 2.7 ± 0.4 |
| #E41G, F50L, I57A | 331.6 ± 0.5 | 191 ± 3 | 1.5 ± 0.1 | 11.5 ± 0.2 | -16.6 ± 0.4 | 1.4 ± 0.0 |
| A16T, N56I | 339.4 ± 0.6 | 227 ± 19 | 4.1 ± 0.3 | 7.0 ± 0.4 | -17.0 ± 1.5 | 2.4 ± 0.2 |
| #V31D, I57A | 344.7 ± 0.5 | 222 ± 0 | 3.7 ± 0.0 | 7.0 ± 0.0 | -17.7 ± 0.1 | 2.5 ± 0.0 |
| #I57A | 333.1 ± 0.5 | 269 ± 14 | 5.4 ± 0.5 | 11.6 ± 0.6 | -17.9 ± 0.6 | 1.5 ± 0.1 |
| G10E, T39S | 339.7 ± 0.7 | 238 ± 16 | 4.0 ± 0.7 | 8.5 ± 0.5 | -18.4 ± 0.4 | 2.2 ± 0.1 |
| #M40T, R48S, I57V | 344.3 ± 0.3 | 235 ± 9 | 3.8 ± 0.2 | 7.1 ± 0.1 | -19.1 ± 0.4 | 2.7 ± 0.1 |
| G35W, A58V | 336.8 ± 0.1 | 281 ± 19 | 5.7 ± 0.5 | 6.9 ± 0.2 | -19.1 ± 0.9 | 2.8 ± 0.1 |
| E4G | 341.5 ± 0.1 | 259 ± 7 | 4.0 ± 0.2 | 7.3 ± 0.1 | -21.4 ± 0.3 | 2.9 ± 0.1 |
| L32Q, I57S | 348.1 ± 0.6 | 275 ± 5 | 4.4 ± 0.0 | 6.1 ± 0.2 | -22.9 ± 0.7 | 3.8 ± 0.2 |
| #V13E | 343.1 ± 0.2 | 280 ± 23 | 4.1 ± 0.5 | 9.6 ± 0.6 | -24.0 ± 1.6 | 2.5 ± 0.2 |
| #E15G, M40T, R48C, I57V | 347.6 ± 0.1 | 281 ± 8 | 4.1 ± 0.1 | 8.3 ± 0.2 | -24.9 ± 0.7 | 3.0 ± 0.1 |
| #V38E, I57V | 348.8 ± 0.3 | 295 ± 16 | 4.4 ± 0.3 | 8.9 ± 0.5 | -25.8 ± 1.2 | 2.9 ± 0.2 |
| T39A, I57N | 343.5 ± 0.2 | 312 ± 0 | 4.7 ± 0.0 | 10.6 ± 0.1 | -26.4 ± 0.1 | 2.5 ± 0.0 |
| L49I | 350.3 ± 0.2 | 322 ± 8 | 4.6 ± 0.1 | 8.7 ± 0.2 | -28.9 ± 0.8 | 3.3 ± 0.1 |
| V34E, D45Y | 344.3 ± 0.0 | 363 ± 7 | 6.0 ± 0.2 | 8.4 ± 0.2 | -29.2 ± 0.5 | 3.5 ± 0.1 |
| **wild-type** | **352.3 ± 0.2** | **322 ± 11** | **4.3 ± 0.2** | **8.2 ± 0.1** | **-30.5 ± 0.8** | **3.7 ± 0.1** |
| P33L | 354.1 ± 0.5 | 330 ± 43 | 4.2 ± 0.7 | 8.3 ± 0.8 | -32.5 ± 3.5 | 3.9 ± 0.6 |
| #L49I, I57V | 356.2 ± 0.3 | 332 ± 13 | 4.2 ± 0.2 | 7.9 ± 0.4 | -32.9 ± 1.3 | 4.2 ± 0.3 |
| #I57V | 354.5 ± 0.2 | 349 ± 12 | 4.7 ± 0.2 | 8.3 ± 0.2 | -33.3 ± 0.9 | 4.0 ± 0.1 |
| #D55G, I57V | 361.6 ± 0.3 | 364 ± 13 | 4.3 ± 0.2 | 8.5 ± 0.2 | -38.1 ± 0.8 | 4.5 ± 0.1 |
| D55G | 361.1 ± 0.1 | 378 ± 10 | 4.7 ± 0.1 | 8.5 ± 0.2 | -38.4 ± 1.0 | 4.5 ± 0.2 |

#Variant selected from the I57A library.

### Comparing with computational stability predictors

To compare how well the stability of the selected set of CI2 variants can be predicted computationally we used FoldX, Rosetta and a sequence-based method that analyses variability in a multiple sequence alignment of homologous sequences (hereafter referred to as SEQ), to calculate ΔΔ*G*_f_ for each variant (Figure 2). The overall performances of these computational methods on our CI2 variants are similar to benchmarks tests on other sets of proteins (Khan and Vihinen, 2010; Potapov et al., 2009). The Pearson’s correlation coefficients (*r*) of the experimentally determined values with those calculated using FoldX, Rosetta and SEQ are 0.81, 0.74 and 0.72, respectively. In general, the three stability predictors agree in the overall effect of mutations, but they are inconsistent in the exact value. Importantly, while the methods are relatively good at predicting destabilizing effects, they are generally not able to predict stabilized variants, though FoldX does predict D55G to be highly stabilizing.

**Figure 2.**
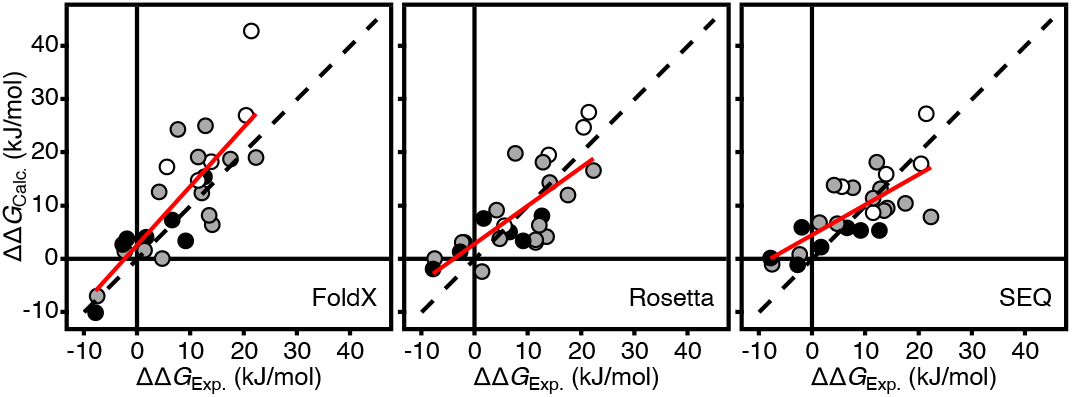
Correlations between experimentally determined and predicted DD*G*_f_ values. The predicted values were calculated by FoldX, Rosetta and SEQ as indicated in the lower right part of each panel. The dotted line shows the identity line and the solid red line is the best fit straight line. Each data point represents one of the variants in Table 1 and the data points are coloured according to the number of amino acid substitutions (black – one substitution; grey – two substitutions; white – three or more substitutions). The scale for the SEQ method is in arbitrary units.

### Analysis of double mutant cycles

In an attempt to understand better the origin of the increased stability of the two double mutants (D55G/I57V and L49I/I57V), we performed a more detailed analysis of the thermodynamic cycles from the wild-type through the single mutants to the double mutants. To compare the variants in the two double mutant cycles we re-analysed the stability data assuming a common *m*-value of 8.4 ± 0.3 kJ/mol/M corresponding to the average of the *m-*values for wild-type, L49I, D55G, I57V, D55G/I57V and L49I/I57V listed in Table 1. As long as there is no significant change in the solvent accessible surface area exposed upon unfolding the *m*-value is also expected not to change (Myers et al., 1995). As done previously (Itzhaki et al., 1995), we have therefore used the average *m*-value for comparing differences in the free energy for folding (ΔΔ*G*_f_). The normalized stability curves originating from this analysis are shown in Figure 3a. Relative to the wild-type, I57V is stabilized by ΔΔ*G*_f_ = −2.2 ± 0.1 kJ/mol (Figure 3b). For D55G ΔΔ*G*_f_ = −6.9 ± 0.3 kJ/mol and combining D55G and I57V leads to no further stabilization (ΔΔ*G*_f_ = −6.5 ± 0.2 kJ/mol). Indeed, the two mutations have an unfavourable synergistic effect, ΔΔΔ*G*_f_, of 2.6 ± 0.4 kJ/mol. In contrast, introducing L49I, which on its own destabilizes by ΔΔ*G*_f_ = 3.4 ± 0.1 kJ/mol, together with I57V results in a total stabilization of the L49I/I57V double mutant of ΔΔ*G*_f_ = −3.8 ± 0.1 kJ/mol. In this case the synergistic effect of introducing both L49I and I57V is ΔΔΔ*G*_f_ −5.1 ± 0.2 kJ/mol.

**Figure 3.**
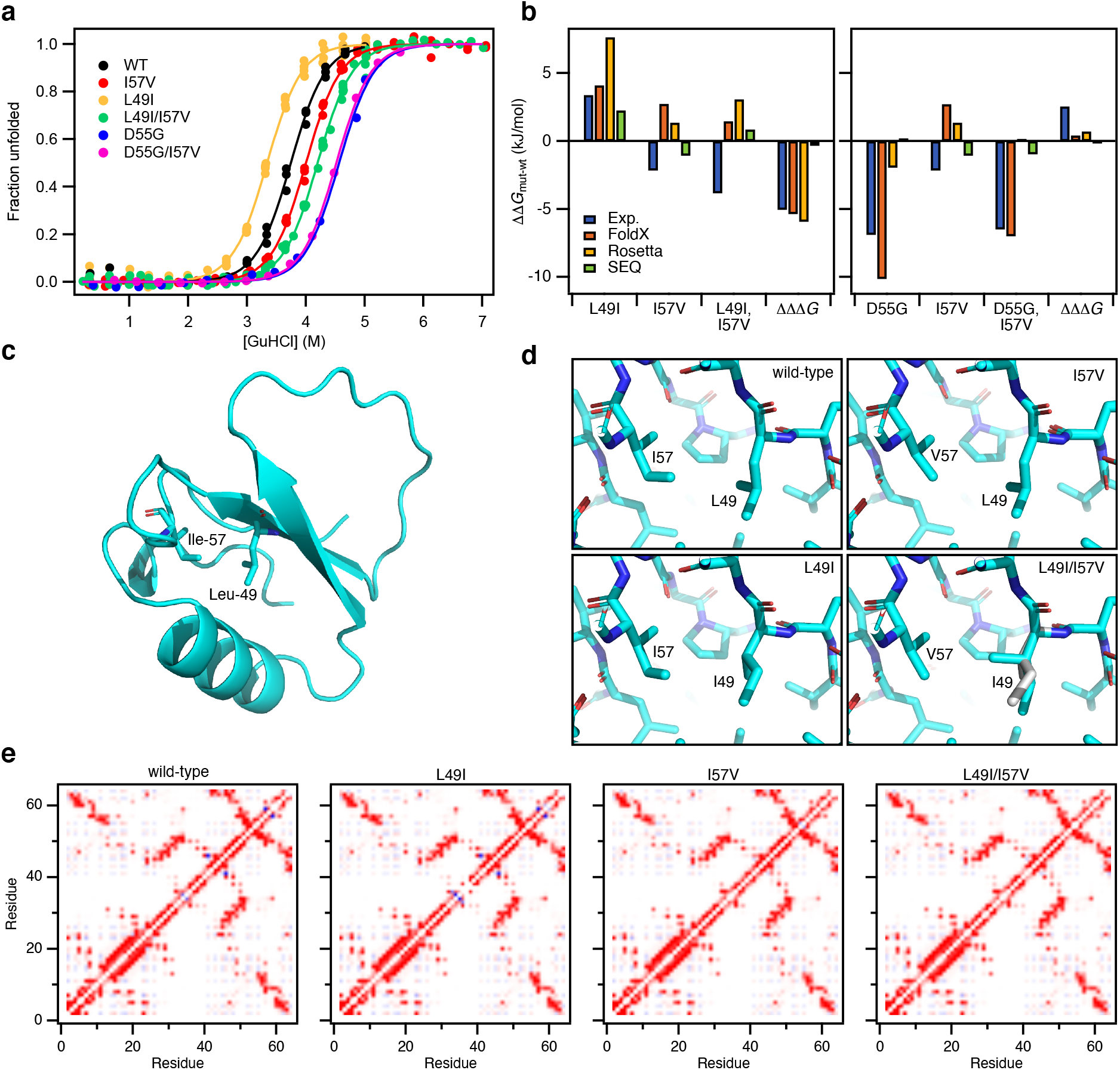
Stability and structural analysis of stabilized CI2 double mutants. (**a**) Equilibrium stability curves of CI2 mutants at 25 °C. The experimental data (filled circles) were fit to a model for two state folding (solid line) keeping the *m*-value fixed at 8.4 kJ/mol/M. The data are here normalized to show the degree of unfolding for direct comparison and visualization only. (**b**) Differences in stability relative to wild-type CI2, ΔΔ*G*_f_, for the variants in the L49I/I57V (left) and D55G/I57V (right) double mutant cycles. Experimental ΔΔ*G*_f_ as well as ΔΔ*G*_f_ predicted by FoldX, Rosetta and SEQ are shown. The ΔΔΔ*G* is the additional contribution to the conformational stability of the double mutants compared to the sum of the contribution of the single mutants, ΔΔ*G*_f,mut1/mut2_ – (ΔΔ*G*_f,mut1_ + ΔΔ*G*_f,mut2_). The energy for the SEQ method is in arbitrary units. (**c**) Overview of the structure of CI2 with the locations of L49 and I57. (**d**) Structural details around positions 49 and 57 in the four CI2 variants in the wild-type to L49I/I57V double mutant cycle. (**e**) Residue wise contact maps. The pairwise interaction energies were calculated from AMBER FF99 using IEM (Bendová-Biedermannová et al., 2008). The colour scale is from dark red (−8 kJ/mol) to dark blue (8 kJ/mol).

To evaluate if the observed effects of the double mutants could have been predicted, we compared with the expected ΔΔ*G*_f_ from FoldX, Rosetta and SEQ. Particularly, FoldX does a good job in predicting the effects of the L49I and D55G mutations (Figure 3b). None of the tools work well for predicting ΔΔ*G*_f_ for I57V or L49I/I57V. FoldX predicts the effect of D55G/I57V rather well, but this can be attributed to the dominating effect of the D55G mutation that was well predicted on its own. It appears as if ΔΔΔ*G*_f_ of L49I/I57V is well predicted by both FoldX and Rosetta (Figure 3b). However, the individual ΔΔ*G*_f_ used for the calculations are not correct and in most cases the signs of the values are incorrect. We thus conclude that the outcome of the double mutations could not have been predicted.

### Structural analysis

The positive synergistic effect of L49I and I57V is interesting to investigate in more detail to gain insight into how proteins may be designed or evolve to become more stable. We therefore determined the crystal structures of all four CI2 variants in the wild-type to L49I/I57V double mutant cycle. Overall the structures of L49I, I57V and L49I/I57V are highly similar to the structure of the wild-type (Figure 3c) with RMSDs for the backbone of 0.16 Å, 0.45 Å and 0.17 Å. The larger RMSD for I57V is a result of the overhand loop in this structure adopting an alternative conformation compared to the other three structures. This conformation is similar to the conformation of the overhand loop in the older crystal structure of wild-type CI2 (PDB-code 2CI2) (McPhalen and James, 1987). Excluding residues 44-50 in this loop from the comparison reduces the backbone RMSD to the wild-type to 0.14 Å, 0.16 Å and 0.15 Å for L49I, I57V and L49I/I57V, respectively.

Around the mutated residues the structural changes are minimal (Figure 3d). The Val at position 57 in I57V and L49I/I57V superimpose, except for the missing δ1 methyl group, with the Ile at position 57 in the wild-type and L49I. The Ile at position 49 in L49I is oriented similarly to the Leu in wild-type and I57V. In the structure of the double mutant, however, we observe that the Ile at position 49 is found in two alternative conformations. The minor conformation, accounting for roughly 30% of the electron density, has a conformation similar to that seen in L49I. In the major conformation that accounts of the remaining 70% of the electron density, the g2 methyl is rotated approximately 120° and points towards the position where the δ1 methyl group of the Ile at position 57 would be in the wild-type structure.

Analysis of the pairwise interaction energies at the residue level (Figure 3e) suggests that much of the stabilizing effect of the I57V mutation originates from an unfavourable interaction between I57 and Q59 that is observed in structures of both the wild-type and L49I. The effect of the L49I mutation, which destabilizes the wild-type, but stabilizes the I57V variant is more subtle and not easily explained from the crystal structures. It appears that some of the destabilization of the L49I mutation is a long-range effect resulting in several less favourable interactions among residues 57-62. These negative effects are relieved in the double mutant (Figure 3e).

## Discussion

Using our bacterial stability sensor, we selected 25 stable and cooperatively folded variants of CI2 from libraries of random mutations. We thus demonstrate that the system can be used as a screening assay for methods like directed evolution and deep mutational scanning to select protein variants with both stabilizing and destabilizing effects originating from mutant libraries. When using the system, it is important that the dynamic range of GFP fluorescence will allow changes in stability to be observed. To select variants with increased stability and thus low GFP fluorescence, it is necessary that starting point for the mutagenesis has a relatively high GFP fluorescence and *vice versa.* Screening assays for higher protein stability are often applied when the starting point has low stability, and in such cases a high initial GFP signal is expected. In the case of CI2, however, it was necessary to introduce the destabilizing I57A mutation to get a proper GFP signal of the background. The Ala was introduced by changing the 57^th^ codon to GCG. To mutate this codon into an Ile codon three base substitutions are needed. It is thus highly unlikely that a revertant to the wild-type will be generated by error prone PCR, and we indeed did not observe the wild-type sequence in this library. Instead the most abundant mutant after selection was I57V which can be made by a single base substitution from the Ala codon. As I57V is more stable than the wild-type this demonstrates the efficiency of the selection system. We note that mutations in the predicted DnaK binding sites of CI2 (Figure 1c) will also lead to decrease in the GFP signal and thus appear similar to mutations that stabilize the protein. This is seen for I30K and V31D selected in the I57A background that are highly unstable. Variants with amino acid substitutions in the DnaK binding site 17-23 could not even be purified, suggesting that the decreased GFP signal originates from decreased interactions with DnaK rather than increased protein stability. Indeed, while both I30K/I57A and I30K/I57V were selected to give decreased GFP signal compared to I57A, only I30K/I57V could successfully be purified. Thus, we expect that some of the variants that could not be purified reflect substitutions with intrinsically weaker DnaK binding in their unfolded states, rather than variants that restabilize I57A. This is supported by the distribution of predicted DnaK binding propensities for the variants that could not be purified that is shifted to lower scores compared to the distribution of those variants that were successfully purified (Figure S2c). We suggest that our sensor system in the future might also be used to screen for peptides that bind DnaK.

As all the variants selected from the I57A background carry either the I57A or the I57V it is not surprising that many double mutants were selected. Two of these (L49I/I57V and D55G/I57V) were highly stabilized, both relative to the I57A background but also more than the wild-type protein. In an attempt to understand the origin of this increased stability we also included the single mutants L49I and D55G in our analysis to make a thermodynamic double mutant cycle. For D55G/I57V the increased stability is almost completely an effect of the D55G mutation. We suggest that is a result of the residue at position 55 being located in the α_L_ region of the Ramachandran map, where Gly is even more common than Asp (Hovmöller et al., 2002). Repulsive interactions with nearby E14 could also play a role. For L49I/I57V on the other hand there is a large non-additive effect. From the crystal structures a few subtle changes in the interaction energies that could contribute to stabilization were identified. However, none of the prediction methods that we used were able to pick up these effects. One explanation for this is that the changes FoldX and Rosetta make to a structure to accommodate a mutation are local. If long range changes are important to explain the change in stability the methods will miss them. Furthermore, errors may accumulate when multiple mutations are introduced (Figure 2).

In conclusion, we have generated a set of variants in CI2 with varying stabilities and most of them containing multiple amino acid substitutions. The data could contribute to optimizing stability predicters. Of particular interest is the synergistic effect of the two substitutions in L49I/I57V. Although we see small structural changes in the structure compared with the other structures in the mutant cycle, the presence of two distinct conformations of the Ile at position 49 in the double mutant could point to effects of conformational entropy or conformational changes also contributing.

## Materials and methods

### CI2 mutant libraries

The wild-type CI2 sequence used here is UniProt: P01053, residues 22-84 with an additional N- terminal Met. The numbering we use, start at this Met, which is the numbering system commonly used in the literature. cDNA encoding this sequence was inserted into a pET22-mCherry vector between the *Nde*I and *Hind*III restriction sites to preserve the translation coupling using Gibson assembly. The QuickChange Lightning II mutagenesis kit (Agilent) was used to create the I57A variant in the CI2_WT_pET22_mCherry vector. Mutant libraries of CI2 WT and CI2 I57A were generated using the GeneMorph II Random mutagenesis kit (Agilent). The mutation frequency was aimed at 0-4 amino acid substitutions per gene by adjusting the initial target DNA and the number of amplification cycles. The PCR product from the random mutagenesis was used as a MEGAprimer for the insertion of the mutants into the CI2 WT or CI2 I57A backgrounds. The PCR products were transformed into MegaX DH10B T1^R^ Electrocomp Cells (Invitrogen) following manufacturer’s instructions and everything was plated on LB agar plates with 100 μg/mL ampicillin.

The CI2 libraries were transformed into electrocompetent Rosetta2(DE3)pLysS pSEVA631-IBpAP- GFP-ASV cells (Zutz et al., 2020). After recovery, the transformants were directly inoculated in 3 mL LB medium containing 100 μg/mL ampicillin, 25 μg/mL chloramphenicol and 50 μg/mL spectinomycin and grown overnight at 37 °C and 180 rpm. Cells were transferred into fresh medium and grown at 30 °C and 250 rpm to an OD_600_ of 0.5 – 0.7. Expression was induced by addition of 0.5 mM IPTG and the growth temperature of the culture was shifted to 30 °C.

1 h after induction cells were analyzed by flow cytometry (Instrument: BD FACS-Aria SORP cell sorter; Laser 1: 488 nm: >50 mW, Filter: 505LP, 515/20-nm FITC; Laser 2: 561 nm: >50 mW; Filter: 600LP, 610/20-nm PE-Texas Red). 100,000 cells expressing a CI2 WT mutant protein with increased GFP signal, and 100,000 cells expressing a CI2 I57A mutant protein with decreased GFP signal were sorted in 2 mL LB medium supplemented with antibiotics and grown overnight at 37 °C and 300 rpm. To further enrich the *E. coli* fraction harboring proteins with altered protein stability, protein expression was induced again and cells (100,000 events) were sorted as described above. The following day, the sorted cell population was analyzed 1 hour after induction of protein expression by flow cytometry (Instrument: BD FACS-Aria SORP cell sorter; Laser 1: 488 nm: >50 mW, Filter: 505LP, 515/20-nm FITC; Laser 2: 561 nm: >50 mW; Filter: 600LP, 610/20-nm PE-Texas Red). Single cells were sorted directly into 100 μl LB supplemented with antibiotics in 96 well culture plates and grown overnight at 37 °C and 300 rpm. The single cells were characterized by Sanger sequencing.

### Library preparation for NGS

Samples from the CI2 WT library were extracted from each step of FACS selection for NGS. 2 mL culture was centrifuged for 10,000 x g for 2 minutes and plasmids purified using the NucleoSpin^®^ Plasmid kit (Machery-Nagel). The CI2 genes were amplified using the Phusion Hot Start II DNA Polymerase (Thermo Scientific) and region of interest-specific primers with overhang adapters. The PCR amplicons were purified using AMPure XP beads (Beckman Coulter) following manufacturer’s instructions. Dual indices and Illumina sequencing adapters (Nexteras XT Index kit (Nextera v2 D)) were attached using the KAPA HiFi HotStart DNA polymerase. The amplicon libraries were purified using AMPure XP beads (Beckman Coulter) following manufacturer’s instructions. Concentrations of the purified amplicon libraries were quantified using a Qubit^®^ 2.0 Fluorometer and the dsDNA broad range kit. To determine the average bp length, a bioanalyzer and the DNA 1000 kit (Agilent) was used following manufacturer’s instructions. The libraries were normalized to 10 nM and all libraries were pooled. The samples were spiked with 5 % Phi-X control DNA (Illumina) and loaded onto the flow cell and sequenced on and then applied onto an Illumina MiSeq instrument.

### Cloning, Expression and stability of single sorted CI2 clones

Glycerol stocks of single cells sorted from the CI2 libraries were used as DNA template in colony PCR using the Phusion Hot Start II High-Fidelity DNA polymerase (Thermo Scientific). CI2 genes were amplified and ligated into a pET11a vector using the *Nde*I and *Bam*HI restriction sites. All clones were verified using Sanger sequencing. All protein expression were performed in BL21(DE3)pLysS. For small scale protein expression, the bacteria were grown in 2 ml ZYM5052 autoinduction media in a 24 well cell culture plate with incubation at 37 °C for 4 hours followed by 20 hours at 20 °C. Low expression level plasmids were expressed in 50 mL ZYM5052 autoinduction media. Cells were harvested by centrifugation at 5000 x g for 15 minutes. The pellet frozen at −20 °C and resuspended in 10 mM Na-acetate, pH 4.4 before centrifugation at 20,000 x g for 30 minutes. The supernatant was further diluted in the same buffer. The samples were applied onto a 1 ml Resource S column equilibrated with 20 mM Na-acetate, pH 4.4 and step eluted with 20 mM Na-acetate, pH 4.4, 1 M NaCl. The the peak fraction was applied onto a Superdex 75 16/85 column equilibrated with 50 mM NH_4_HCO_3_, and the fractions containing CI2 were collected and lyophilized before dissolving, in 50 mM MES, pH 6.25. For expression of protein for structure determination the bacteria were grown in LB media. The expression was induced by 0.4 mM IPTG at OD_600_ 0.6 – 0.8. Cells were harvested by centrifugation at 5000 x g for 15 min and resuspended in 25 mM Tris- HCl, pH 8, 1 mM EDTA before lysis by two freeze-thaw cycles. The sample was cleared by centrifugation at 20,000 x g at 4 °C. Polyetylenimine was added to a concentration of 1 % and the sample centrifuged for 15 min at 20,000 x g. Ammonium sulphate was added to the supernatant to 70 % saturation, and left for 30 min at 4 °C before centrifugation at 20,000 x g. The pellet was resuspended in 25 mM Tris-HCl, pH 8 and heated at 40°C until all precipitate was solubilized. The samples were centrifuged at 20,000 x g for 10 min before size exclusion chromatography in 10 mM NH_4_HCO_3_. Peak fractions were pooled and lyophilized and finally resuspended in MilliQ water and dialysed against water.

### Equilibrium unfolding

Protein concentrations were determined by absorbance at 280 nm measured on a NanoDrop 1000 due to the low volume samples. Equilibrium stability in GuHCl was measured with a final protein concentration of 10 μM at 13-16 concentrations of GuHCl evenly distributed in the range from 0 to 5, 6 or 7 M depending on the stability of the variant. The degree of unfolding was followed by fluorescence measurements on a Prometheus NT.48 (nanoTemper technologies) using Prometheus NT.48 high sensitivity capillaries. The temperature was ramped from 15 to 95°C with a temperature increment of 1°C/min. Global analysis of temperature and solvent denaturation was performed as described (Hamborg et al., 2020).

### Computational prediction of stability and DnaK binding

Version 4 of the FoldX energy function (Schymkowitz et al., 2005) was used to estimate the free- energy change upon mutations of CI2, using the coordinates of the PDB entry 2CI2 (McPhalen and James, 1987). The RepairPDB function of FoldX was first applied to the wild type structure. The resulting structure was used as input to the BuildModel function to generate the models of the investigated mutants and to evaluate their ΔΔ*G*_f_.

The Rosetta energy function (Kellogg et al., 2010) in its cartesian version (Park et al., 2016) was also used to estimate ΔΔ*G*_f_, using the coordinates of PDB entry 2CI2. The wild-type structure was first relaxed in cartesian space with restrained backbone and sidechain coordinates. The resulting coordinates were then used to build the model of the investigated mutants and to evaluate their ΔΔ*G*_f_ by means of the Cartesian_ddg function. The calculations were repeated on five independent runs, whose results were then averaged to obtain the final values reported in the manuscript. The resulting difference in stability was multiplied by 1.44 to bring the ΔΔG values from Rosetta energy units onto a scale corresponding to kJ/mol (Jepsen et al., 2020).

Looking at the mutational pattern observed in a multiple sequence alignment of homologous sequence, it is possible to build a global statistical model of the relative protein family variability (Marks et al., 2011), which takes into account not only single-site conservation, but also correlated mutations between site pairs. This approach aims at exploiting the structural and functional constraints encoded in the family evolution (Granata et al., 2017), assigning to each specific sequence a score related to the probability of being a good representative of that family (Figliuzzi et al., 2015). Even if this measure is more related to the general fitness of the sequence, it can also be used *bona fide* to judge the effect of a specific mutation on protein stability. In order to have a variation model which is statistically significant, we obtained a larger multiple sequence alignment containing CI2 homologues by building a hidden Markov model of the protein family, based on 4 iterations of Jackhmmer (Finn et al., 2015) and extracting the sequences from the Uniprot Uniref100 database (Suzek et al., 2014). The sequence containing more than 50% of gaps with respect to wild-type sequence were excluded, together with the sequences sharing more than 90% of sequence identity, resulting in an alignment of 942 independent sequences. We then used the asymmetric plmDCA algorithm (Ekeberg et al., 2013) to calculate the parameters of the sequence model. The score of the wild-type sequence was then subtracted to the one of each analysed sequence to obtain the final values reported in the manuscript.

We used the Limbo algorithm (Durme et al., 2009) to predict DnaK binding sites in both wild-type and variant forms of CI2.

### Crystallization, diffraction experiments and structure calculations

CI2 WT, L49I and L49I/I57V were crystalized at 293 K in 40% (NH_4_)_2_SO_4_, 50 mM Tris-HCl, pH 8.0 at a protein concentration of 75 mg/ml. CI2 I57V were crystalized in 0.1 M Tris-HCl, 8% PEG 8000, pH 8.5. Data for WT were collected on an inhouse setup with an Agilent SuperNova diffraction source (1.5406 Å) and an Atlas CCD detector. Data for L49I, I57V and L49I/I57V were collected at DESY, Hamburg, beamline P13 (0.9763 Å) equipped with a Pilatus 6M-F, S/N 60-0117-F detector.

The reflections were collected using autoPROC (Vonrhein et al., 2011), which also scales and merges the data using the CCP4 programs Pointless and Aimless (Winn et al., 2011) as well as Staraniso (http://staraniso.globalphasing.org/cgi-bin/staraniso.cgi). The latter program was employed because of pronounced anisotropic distribution of reflections. The structures were solved by molecular replacement using the CCP4 pogram Phaser (McCoy et al., 2007). The initial search model was wild-type CI2 (PDB code 2CI2). Later molecular replacement solutions were obtained using the higher resolution structures described in the present paper. The structures were carefully examined, adjusted and refined with Coot and Refmac5, respectively (Emsley et al., 2010; Murshudov et al., 2011). To make sure that the structures were not in a domain swapped configuration (Campos et al., 2019), molecular replacement solutions were also sought using the domain swapped structure with PDB accession code 6QIZ. These consistently yielded significantly worse statistics.

## Acknowledgements

This work was supported by the Novo Nordisk Foundation [grant numbers NNF15OC0016360 and NNF18OC0033926]. The authors thank Pia Skovgaard for technical assistance. KLL and KT are members of Integrative Structural Biology at the University of Copenhagen (www.isbuc.ku.dk). We thank Profs. F. Rousseau and J. Schymkowitz (VIB) for sharing the Limbo Software. We acknowledge excellent support at the P14 beamline operated by EMBL Hamburg at the PETRA III storage ring (DESY, Hamburg). We are grateful for support from the DANSCATT program of the Danish Council for Research and Innovation. We thank Thomas Lykke-Møller Sørensen (Aarhus University) for help with the data acquisition at DESY. Finally, we thank Pernille Harris for access to the in-house diffractometer at the Technical University of Denmark.

## Data availability

The atomic coordinates and structure factors for the structures presented in this work are deposited at the Protein Data Bank (https://www.wwPDB.org) under accession numbers 7A1H, 7A3M, 7AOK and 7AON. Other data included in the figures and that support the findings of this study are available from the corresponding author upon reasonable request.

## References

Bendová-Biedermannová L, Hobza P, Vondrášek J. 2008. Identifying stabilizing key residues in proteins using interresidue interaction energy matrix: Pair-Wise Interaction Energy Matrix. Proteins Struct Funct Bioinform 72:402–413. doi:10.1002/prot.21938

Bershtein S, Segal M, Bekerman R, Tokuriki N, Tawfik DS. 2006. Robustness–epistasis link shapes the fitness landscape of a randomly drifting protein. Nature 444:929–932. doi:10.1038/nature05385

Campos LA, Sharma R, Alvira S, Ruiz FM, Ibarra-Molero B, Sadqi M, Alfonso C, Rivas G, Sanchez-Ruiz JM, Garrido AR, Valpuesta JM, Muñoz V. 2019. Engineering protein assemblies with allosteric control via monomer fold-switching. Nat Commun, Nature Communications 10:5703. doi:10.1038/s41467-019-13686-1

Capriotti E, Fariselli P, Casadio R. 2005. I-Mutant2.0: predicting stability changes upon mutation from the protein sequence or structure. Nucleic Acids Res 33:W306–W310. doi:10.1093/nar/gki375

Dehouck Y, Grosfils A, Folch B, Gilis D, Bogaerts P, Rooman M. 2009. Fast and accurate predictions of protein stability changes upon mutations using statistical potentials and neural networks: PoPMuSiC-2.0. Bioinformatics 25:2537–2543. doi:10.1093/bioinformatics/btp445

Durme JV, Maurer-Stroh S, Gallardo R, Wilkinson H, Rousseau F, Schymkowitz J. 2009. Accurate prediction of DnaK-peptide binding via homology modelling and experimental data. PLoS Comput Biol 5:e1000475. doi:10.1371/journal.pcbi.1000475

Ekeberg M, Lövkvist C, Lan Y, Weigt M, Aurell E. 2013. Improved contact prediction in proteins: Using pseudolikelihoods to infer Potts models. Phys Rev E Stat Nonlin Soft Matter Phys 87:012707. doi:10.1103/physreve.87.012707

Emsley P, Lohkamp B, Scott WG, Cowtan K. 2010. Features and development of Coot. Acta Crystallogr Sect D Biol Crystallogr 66:486–501. doi:10.1107/s0907444910007493

Figliuzzi M, Jacquier H, Schug A, Tenaillon O, Weigt M. 2015. Coevolutionary Landscape Inference and the Context-Dependence of Mutations in Beta-Lactamase TEM-1. Mol Biol Evol 33:268–80. doi:10.1093/molbev/msv211

Finn RD, Clements J, Arndt W, Miller BL, Wheeler TJ, Schreiber F, Bateman A, Eddy SR. 2015. HMMER web server: 2015 update. Nucleic Acids Res 43:W30–W38. doi:10.1093/nar/gkv397

Foit L, Morgan GJ, Kern MJ, Steimer LR, Hacht AA von, Titchmarsh J, Warriner SL, Radford SE, Bardwell JCA. 2009. Optimizing Protein Stability In Vivo. Mol Cell, Molecular Cell 36:861–871. doi:10.1016/j.molcel.2009.11.022

Goldenzweig A, Goldsmith M, Hill SE, Gertman O, Laurino P, Ashani Y, Dym O, Unger T, Albeck S, Prilusky J, Lieberman RL, Aharoni A, Silman I, Sussman JL, Tawfik DS, Fleishman SJ. 2018. Automated Structure- and Sequence-Based Design of Proteins for High Bacterial Expression and Stability. Mol Cell 70:380. doi:10.1016/j.molcel.2018.03.035

Granata D, Ponzoni L, Micheletti C, Carnevale V. 2017. Patterns of coevolving amino acids unveil structural and dynamical domains. Proc Natl Acad Sci USA 114:E10612–E10621. doi:10.1073/pnas.1712021114

Gromiha MM, Anoosha P, Huang L-T. 2016. Applications of Protein Thermodynamic Database for Understanding Protein Mutant Stability and Designing Stable Mutants. Methods Mol Biol 1415:71–89. doi:10.1007/978-1-4939-3572-7_4

Guerois R, Nielsen JE, Serrano L. 2002. Predicting Changes in the Stability of Proteins and Protein Complexes: A Study of More Than 1000 Mutations. J Mol Biol, Journal of Molecular Biology 320:369–387. doi:10.1016/s0022-2836(02)00442-4

Hamborg L, Horsted EW, Johansson KE, Willemoës M, Lindorff-Larsen K, Teilum K. 2020. Global analysis of protein stability by temperature and chemical denaturation. Anal Biochem 605:113863. doi:10.1016/j.ab.2020.113863

Hovmöller S, Zhou T, Ohlson T. 2002. Conformations of amino acids in proteins. Acta Crystallogr Sect D Biol Crystallogr 58:768–776. doi:10.1107/s0907444902003359

Itzhaki LS, Otzen DE, Fersht AR. 1995. The Structure of the Transition State for Folding of Chymotrypsin Inhibitor 2 Analysed by Protein Engineering Methods: Evidence for a Nucleation-condensation Mechanism for Protein Folding. J Mol Biol, Journal of molecular biology 254:260–288. doi:10.1006/jmbi.1995.0616

Jackson SE, Fersht AR. 1991a. Folding of chymotrypsin inhibitor 2. 1. Evidence for a two-state transition. Biochemistry, Biochemistry 30:10428–35.

Jackson SE, Fersht AR. 1991b. Folding of chymotrypsin inhibitor 2. 2. Influence of proline isomerization on the folding kinetics and thermodynamic characterization of the transition state of folding. Biochemistry, Biochemistry 30:10436–43.

Jackson SE, Moracci M, elMasry N, Johnson CM, Fersht AR. 1993. Effect of cavity-creating mutations in the hydrophobic core of chymotrypsin inhibitor 2. Biochemistry, Biochemistry 32:11259–69.

Jepsen MM, Fowler DM, Hartmann-Petersen R, Stein A, Lindorff-Larsen K. 2020. Classifying disease-associated variants using measures of protein activity and stability In: Pey AL, editor. Protein Homeostasis Diseases. Academic Press. pp. 91–107. doi:10.1016/b978-0-12-819132-3.00005-1

Jochens H, Aerts D, Bornscheuer UT. 2010. Thermostabilization of an esterase by alignment-guided focussed directed evolution. Protein Eng Des Sel 23:903–909. doi:10.1093/protein/gzq071

Kaufmann KW, Lemmon GH, DeLuca SL, Sheehan JH, Meiler J. 2010. Practically Useful: What the Rosetta Protein Modeling Suite Can Do for You. Biochemistry 49:2987–2998. doi:10.1021/bi902153g

Kellogg EH, Leaver-Fay A, Baker D. 2010. Role of conformational sampling in computing mutation-induced changes in protein structure and stability: Conformational Sampling in Computing Mutation-Induced Changes. Proteins Struct Funct Bioinform 79:830–838. doi:10.1002/prot.22921

Khan S, Vihinen M. 2010. Performance of protein stability predictors. Hum Mutat 31:675–684. doi:10.1002/humu.21242

Lawrence C, Kuge J, Ahmad K, Plaxco KW. 2010. Investigation of an anomalously accelerating substitution in the folding of a prototypical two-state protein. J Mol Biol, Journal of Molecular Biology 403:446–58. doi:10.1016/j.jmb.2010.08.049

Marks DS, Colwell LJ, Sheridan R, Hopf TA, Pagnani A, Zecchina R, Sander C. 2011. Protein 3D Structure Computed from Evolutionary Sequence Variation. PLoS ONE, PloS one 6:e28766. doi:10.1371/journal.pone.0028766

McCoy AJ, Grosse-Kunstleve RW, Adams PD, Winn MD, Storoni LC, Read RJ. 2007. Phaser crystallographic software. J Appl Crystallogr 40:658–674. doi:10.1107/s0021889807021206

McPhalen CA, James MN. 1987. Crystal and molecular structure of the serine proteinase inhibitor CI-2 from barley seeds. Biochemistry, Biochemistry 26:261–9.

Modarres HP, Mofrad MR, Sanati-Nezhad A. 2016. Protein thermostability engineering. RSC Adv 6:115252–115270. doi:10.1039/c6ra16992a

Murshudov GN, Skubák P, Lebedev AA, Pannu NS, Steiner RA, Nicholls RA, Winn MD, Long F, Vagin AA. 2011. REFMAC5 for the refinement of macromolecular crystal structures. Acta Crystallogr Sect D Biol Crystallogr 67:355–67. doi:10.1107/s0907444911001314

Myers JK, Pace CN, Scholtz JM. 1995. Denaturant m values and heat capacity changes: Relation to changes in accessible surface areas of protein unfolding. Protein Sci, Protein science: a publication of the Protein Society 4:2138–2148. doi:10.1002/pro.5560041020

Neira JL, Itzhaki LS, Ladurner AG, Davis B, Gay G de P, Fersht AR. 1997. Following co-operative formation of secondary and tertiary structure in a single protein module. J Mol Biol 268:185–197. doi:10.1006/jmbi.1997.0932

Pandurangan AP, Ochoa-Montaño B, Ascher DB, Blundell TL. 2017. SDM: a server for predicting effects of mutations on protein stability. Nucleic Acids Res 45:W229–W235. doi:10.1093/nar/gkx439

Park H, Bradley P, Greisen P, Liu Y, Mulligan VK, Kim DE, Baker D, DiMaio F. 2016. Simultaneous Optimization of Biomolecular Energy Functions on Features from Small Molecules and Macromolecules. J Chem Theory Comput 12:6201–6212. doi:10.1021/acs.jctc.6b00819

Parthiban V, Gromiha MM, Schomburg D. 2006. CUPSAT: prediction of protein stability upon point mutations. Nucleic Acids Res 34:W239–W242. doi:10.1093/nar/gkl190

Potapov V, Cohen M, Schreiber G. 2009. Assessing computational methods for predicting protein stability upon mutation: good on average but not in the details. Protein Eng Des Sel, Protein engineering, design & selection: PEDS 22:553–560. doi:10.1093/protein/gzp030

Reetz MT, Carballeira JD, Vogel A. 2006. Iterative Saturation Mutagenesis on the Basis of B Factors as a Strategy for Increasing Protein Thermostability. Angew Chem Int Ed Engl 45:7745–7751. doi:10.1002/anie.200602795

Rohl CA, Strauss CEM, Misura KMS, Baker D. 2004. Protein structure prediction using Rosetta. Methods Enzymol 383:66–93. doi:10.1016/s0076-6879(04)83004-0

Sarkisyan KS, Bolotin DA, Meer MV, Usmanova DR, Mishin AS, Sharonov GV, Ivankov DN, Bozhanova NG, Baranov MS, Soylemez O, Bogatyreva NS, Vlasov PK, Egorov ES, Logacheva MD, Kondrashov AS, Chudakov DM, Putintseva EV, Mamedov IZ, Tawfik DS, Lukyanov KA, Kondrashov FA. 2016. Local fitness landscape of the green fluorescent protein. Nature 533:397–401. doi:10.1038/nature17995

Schymkowitz J, Borg J, Stricher F, Nys R, Rousseau F, Serrano L. 2005. The FoldX web server: an online force field. Nucleic Acids Res 33:W382–W388. doi:10.1093/nar/gki387

Stein A, Fowler DM, Hartmann-Petersen R, Lindorff-Larsen K. 2019. Biophysical and Mechanistic Models for Disease-Causing Protein Variants. Trends Biochem Sci 44:575–588. doi:10.1016/j.tibs.2019.01.003

Steipe B, Schiller B, Plückthun A, Steinbacher S. 1994. Sequence Statistics Reliably Predict Stabilizing Mutations in a Protein Domain. J Mol Biol 240:188–192. doi:10.1006/jmbi.1994.1434

Sullivan BJ, Nguyen T, Durani V, Mathur D, Rojas S, Thomas M, Syu T, Magliery TJ. 2012. Stabilizing Proteins from Sequence Statistics: The Interplay of Conservation and Correlation in Triosephosphate Isomerase Stability. J Mol Biol 420:384–399. doi:10.1016/j.jmb.2012.04.025

Suzek BE, Wang Y, Huang H, McGarvey PB, Wu CH, Consortium U. 2014. UniRef clusters: a comprehensive and scalable alternative for improving sequence similarity searches. Bioinformatics 31:926–32. doi:10.1093/bioinformatics/btu739

Trudeau DL, Lee TM, Arnold FH. 2014. Engineered thermostable fungal cellulases exhibit efficient synergistic cellulose hydrolysis at elevated temperatures. Biotechnol Bioeng 111:2390–2397. doi:10.1002/bit.25308

Vonrhein C, Flensburg C, Keller P, Sharff A, Smart O, Paciorek W, Womack T, Bricogne G. 2011. Data processing and analysis with the autoPROC toolbox. Acta Crystallogr D Biol Crystallogr 67:293–302. doi:10.1107/s0907444911007773

Wijma HJ, Floor RJ, Jekel PA, Baker D, Marrink SJ, Janssen DB. 2014. Computationally designed libraries for rapid enzyme stabilization. Protein Eng Des Sel 27:49–58. doi:10.1093/protein/gzt061

Winn MD, Ballard CC, Cowtan KD, Dodson EJ, Emsley P, Evans PR, Keegan RM, Krissinel EB, Leslie AGW, McCoy A, McNicholas SJ, Murshudov GN, Pannu NS, Potterton EA, Powell HR, Read RJ, Vagin A, Wilson KS. 2011. Overview of the CCP4 suite and current developments. Acta Crystallogr D Biol Crystallogr, Acta crystallographica Section D, Biological crystallography 67:235–242. doi:10.1107/s0907444910045749

Yamashiro K, Yokobori S-I, Koikeda S, Yamagishi A. 2010. Improvement of Bacillus circulans β-amylase activity attained using the ancestral mutation method. Protein Eng Des Sel 23:519–528. doi:10.1093/protein/gzq021

Zutz A, Hamborg L, Pedersen LE, Kassem MM, Papaleo E, Koza A, Herrgård MJ, Teilum K, Lindorff-Larsen K, Nielsen AT. 2020. A dual-reporter system for investigating and optimizing protein translation and folding in E. coli. bioRxiv 2020.09.18.303453. doi:10.1101/2020.09.18.303453

