## Supplementary material for "Synergistic stabilization of a double mutant in CI2 from an *in-cell* library screen": Figure S

**Table S1.** Crystallographic statistics

| *Data set* | **CI2 WT** | **CI2 I57V** | **CI2 L49I** | **CI2 L49I,I57V** |
|---|---|---|---|---|
| PDB code | 7A1H | 7A3M | 7AOK | 7AON |
| **Data collection** | | | | |
| Beamline | N.A. | DESY, B13 | DESY, B13 | DESY, B13 |
| Wavelength (Å) | 1.5406 | 0.97625 | 0.97625 | 0.97625 |
| Temperature (K) | 150 | 100 | 100 | 100 |
| Space group | P622 | P622 | P622 | P622 |
| a, b, c (Å) | 68.27, 68.27, 52.60 | 68.44, 68.44, 52.10 | 68.47, 68.47, 52.94 | 68.39, 68.39, 52.66 |
| α,β,γ (°) | 90, 90, 120 | 90, 90, 120 | 90, 90, 120 | 90, 90, 120 |
| Protein molecules in asymmetric unit | 1 | 1 | 1 | 1 |
| Resolution range (Å) | 15.0 – 1.90 | 59.3 – 1.01 | 59.3 – 1.87 | 59.2 – 1.26 |
| Number of reflections collected | 52781 | 968735 | 183674 | 571649 |
| Number of unique reflections | 6063 | 29927 | 5666 | 15866 |
| Multiplicity | 8.7 (5.2) | 32.4 (5.9) | 32.3 (36.4) | 36.0 (34.1) |
| Completeness (%) | 99.7 (99.7) | 95.4 (60.7) | 88.0 (100.0) | 94.7 (63.4) |
| $R_{\mathrm{pim}}$ | 0.033 (0.332) | 0.015 (0.506) | 0.019 (0.325) | 0.016 (0.462) |
| $CC_{1/2}$ | 0.999 (0.565) | 0.9995 (0.4978) | 0.998 (0.867) | 0.9995 (0.7034) |
| $\langle I/\sigma(I)\rangle$ | 10.4 (1.7) | 22.9 (1.2) | 22.4 (2.5) | 22.6 (1.5) |
| **Refinement statistics** | | | | |
| $R_{\mathrm{work}}/R_{\mathrm{free}}$ (%) | 20.5/25.2 | 14.9/18.1 | 22.0/24.6 | 16.2/21.9 |
| Number of atoms | 551 | 625 | 538 | 598 |
| Water molecules | 28 | 98 | 18 | 60 |
| $\langle B\rangle$ (Å$^2$) | 23.4 | 14.9 | 37.2 | 24.2 |
| Minimal estimated coordinate errors (Å) | 0.123 | 0.003 | 0.015 | 0.005 |
| *RMS deviations from ideal geometry* | | | | |
| Bond lengths (Å) | 0.020 | 0.031 | 0.020 | 0.030 |
| Bond angles (°) | 2.078 | 2.516 | 2.065 | 3.143 |
| Chiral volume (Å$^3$) | 0.111 | 0.140 | 0.109 | 0.192 |
| *Ramachandran plot* | | | | |
| Preferred regions (%) | 98.4 | 98.3 | 96.8 | 96.7 |
| Allowed regions (%) | 1.6 | 1.7 | 3.2 | 3.3 |
| Outliers (%) | 0 | 0 | 0 | 0 |

Numbers in parenthesis refer to outer shell data.

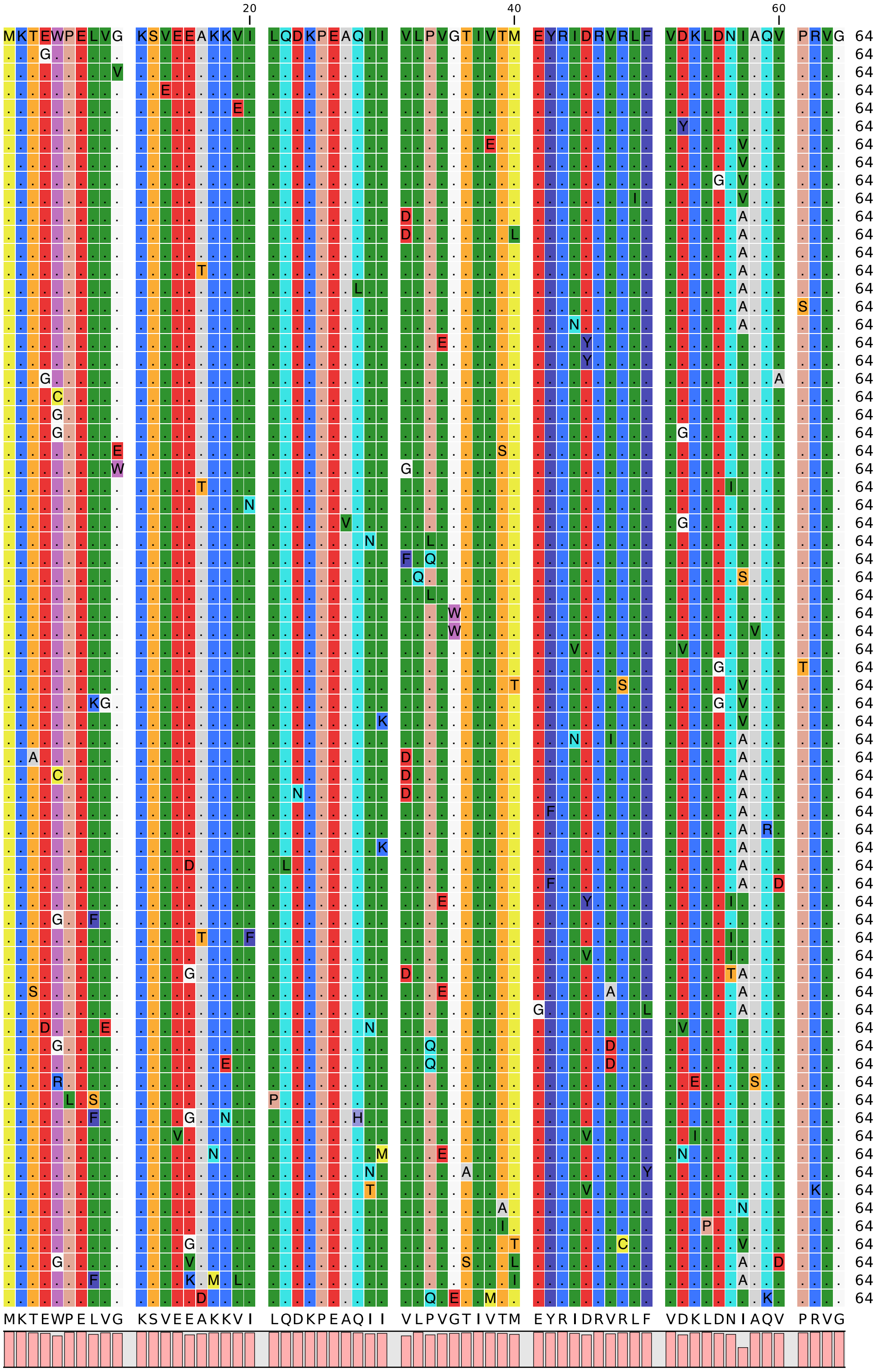

**Figure S1** Alignment of 71 CI2 sequences selected using the folding sensor. The wild-type sequence is shown in the top row. In other rows only amino acid residues that differ from the wild-type sequence are shown as letters. Below the alignment, the consensus sequence is shown on top of bars that corresponds to the level of conservation. Colours indicate amino acid types.

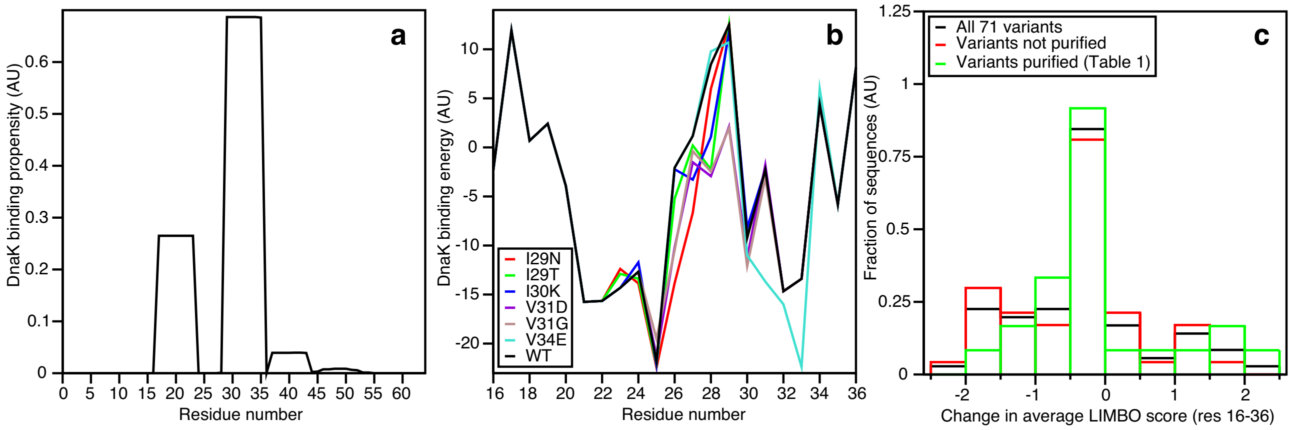

**Figure S2** Propensity for DnaK binding to CI2 as predicted by the Limbo algorithm.[33] (**a**) Sequence dependent DnaK propensity score for wild-type CI2. (**b**) Variation in DnaK binding energy for single point mutations in the major DnaK binding region of CI2. (**c**) Distributions of changes in DnaK binding scores of CI2 variants relative to the score of wild-type CI2, for all 71 initially selected variants (black), those variants that failed to be purified (red), and those that were purified and used for stability measurements (green).
